# Deletion of a KSF Motif Attenuates NSP1 Host Cell Translation Shutoff and Impairs SARS-CoV-2 Virulence

**DOI:** 10.1101/2025.04.16.649178

**Authors:** Chengjin Ye, Shahrzad Ezzatpour, Brian Imbiakha, Nathaniel Jackson, Tolga Cagatay, Anastasija Cupic, Lisa Miorin, Annette Choi, David W. Buchholz, Jordan Carter, Julie Sahler, Ximena A. Olarte-Castillo, Mason C. Jager, Gary R. Whittaker, Avery August, Beatriz Fontoura, Adolfo García-Sastre, Luis Martinez-Sobrido, Hector C. Aguilar

## Abstract

Severe acute respiratory syndrome coronavirus 2 (SARS-CoV-2), the causative agent of coronavirus disease 2019 (COVID-19), triggered a global pandemic with profound social and economic consequences. The viral spike (S) protein has been identified as a key determinant of SARS-CoV-2 pathogenicity. In this study, we demonstrate that the Omicron BA.4 and BA.5 variants, which have closely related S proteins, exhibit different virulence in K18-hACE2 transgenic mice. A comparison of genomic sequences revealed key differences between variants BA.4 and BA.5, including a three amino acid deletion (ΔKSF) in the linker region of the non-structural protein 1 (NSP1) in BA.4. Using reverse genetic systems, we engineered a recombinant (r)SARS-CoV-2 BA.5 expressing BA.4 NSP1, which was significantly attenuated *in vivo*, similar to the natural BA.4 isolate, compared to rBA.5 wild-type (WT). This finding indicates that NSP1 is responsible, at least in part, for the differences in virulence between BA.4 and BA.5. Mechanistically, BA.4 NSP1 showed a reduced ability to inhibit host gene translation compared to BA.5 NSP1. Notably, a rSARS-CoV-2 WA1 original strain containing the same ΔKSF in NSP1 was also attenuated *in vivo* compared to rWA1 WT. Together, these findings highlight the contributions of the NSP1 linker region to inhibiting host gene expression and SARS-CoV-2 pathogenicity, as well as the feasibility of targeting NSP1 for the rational design of live-attenuated vaccines and/or antivirals.

**IMPORTANCE:** Understanding why some SARS-CoV-2 variants cause more severe disease than others is crucial for proper management or for the rational design of prophylactic and therapeutic treatments. While most studies focus on the spike (S) protein, we found that another viral protein, NSP1, also plays a key role in disease severity among SARS-CoV-2 variants. A small deletion in NSP1, present in the Omicron BA.4 variant, weakens the virus ability to shut down the host’s immune response, making BA.4 less severe than BA.5. When we introduced the same deletion into the original SARS- CoV-2 WA1 strain, the virus also became less harmful. This discovery suggests that NSP1 is an important virulence factor and supports the feasibility of targeting NSP1 for the development of new prophylactic and therapeutic treatments against SARS-CoV-2. By uncovering NSP1’s role in pathogenesis, our study provides insights that could help in designing better strategies to combat future variants of SARS-CoV-2.

## INTRODUCTION

Coronaviruses, which cause mild to lethal respiratory infections, are common pathogens of humans and animals (1, 2). Four coronaviruses are endemic in humans (human coronavirus NL63 [HCoV-NL63], HCoV-229E, HCoV-OC43, and HCoV-HKU1) and typically infect the upper respiratory tract, causing common cold symptoms (3). However, three zoonotic coronaviruses (severe acute respiratory syndrome coronavirus [SARS-CoV], Middle East respiratory syndrome coronavirus [MERS-CoV], and severe acute respiratory syndrome coronavirus 2 [SARS-CoV-2]) have been responsible for severe disease in humans over the past two decades (4–6).

The etiological agent of the worldwide Coronavirus Disease 2019 (COVID-19) pandemic, SARS-CoV-2, was first identified in Wuhan, China, in late 2019 in a cluster of patients with pneumonia (7). SARS-CoV-2 belongs to the *Coronaviridae* family of the *Nidovirales* order and is an enveloped positive-sense, single-stranded RNA virus with a non-segmented genome of ∼30 kb in length. SARS-CoV-2 encodes a set of structural (membrane [M], nucleocapsid [N], envelope [E], and spike [S]), non-structural (NSP1-16), and accessory open reading frame (ORF) 1a, 1b, 3a, 6, 7a, 7b, 8, 9b and 10) proteins (8). While structural proteins either hold the RNA genome or are built into the viral particle, NSPs are expressed in infected cells but not incorporated into the virion. In SARS-CoV- 2, the genomic RNA is translated to produce NSPs from two ORFs, 1a and 1b. ORF1a produces polypeptide 1a (pp1a, 440-500 kDa), and the -1 ribosomal frameshift occurs immediately upstream of the ORF1a stop codon, which allows continued translation of ORF1b, yielding a large polypeptide (pp1ab, 740-810 kDa). Sixteen NSPs are co-translationally and post-translationally released from pp1a (NSP1-11) and pp1ab (NSP1-10, NSP12-16) upon proteolytic cleavage by two cysteine proteases encoded by NSP3 (papain-like protease; PLpro) and NSP5 (chymotrypsin-like protease, the main protease, or Mpro) (9).

After four successive COVID-19 pandemic waves driven by Alpha (B.1.1.7), Beta (B.1.351), Gamma (P.1), and Delta (B.1.617.2) variants, Omicron variants emerged in late 2021 and since then have been the dominant circulating strains (10). SARS-CoV-2 Omicron was first documented in South Africa, Botswana, and in a traveler from South Africa in Hong Kong (11). SARS-CoV-2 Omicron variants are more immune evasive but less virulent than previous viral variants (12–14). Omicron variants BA.1 and BA.2 are capable of replicating in K18-hACE2 transgenic mice but result in little to no observable disease or death (15). BA.4 and BA.5 were found to have similar infectivity and pathogenicity than BA.2 in wild-type (WT) and ACE2-expressing hamsters (16). In contrast, our recent findings demonstrated that infection with Omicron BA.5 in K18-hACE2 mice led to marked weight loss, elevated lung pathology, and increased infiltration of inflammatory cells and cytokines in the bronchoalveolar lavage fluid. Notably, BA.5 infection resulted in 100% mortality among 5- to 8-month-old K18-hACE2 mice, a stark difference compared to the outcomes observed with Omicron BA.1 (17). These differences in viral pathogenicity between Omicron BA.1 and BA.5 in K18-hACE2 mice lead us to evaluate whether infection with Omicron BA.4, which is more closely related to BA.5, would lead to the same outcome in K18-hACE2 mice and whether there are viral components contributing to the differences in viral pathogenicity.

Our results suggest that in contrast to Omicron BA.5, BA.4 is attenuated in K18-hACE2 mice, similar to Omicron BA.1. To determine the viral factors responsible for the differences in viral pathogenicity between variants BA.4 and BA.5, we developed an Omicron BA.5 reverse genetic system (RGS). Using this BA.5 RGS, we introduced the NSP1 KSF deletion between amino acids 141 and 143 (ΔKSF), presented in BA.4, into the BA.5 genome. Contrary to the recombinant (r)BA.5, the rBA.5 (ΔKSF) was significantly attenuated in K18-hACE2 mice, similar to BA.4. Notably, infection of K18-hACE2 mice with an rWA1 original strain containing the same NSP1 KSF deletion resulted in significant attenuation compared to its WT counterpart. Mechanistically, the KSF deletion in NSP1 reduced its ability to inhibit host gene expression. These results illustrate that Omicron BA.4 and BA.5, which contain closely related S glycoproteins, exhibit different virulence *in vivo*, demonstrating that viral proteins other than S are important virulence determinants of SARS-CoV-2. These results indicate that in addition to a contribution of the ORF6 D61L mutation present in BA.4 and not in BA.5 (18), the BA. 4-specific NSP1 KSF deletion also has an impact in decreasing virulence. This opens the feasibility of targeting NSP1 for the rational development of attenuation strategies to generate live-attenuated vaccines or for the design of antivirals targeting this important viral virulence determinant.

## RESULTS

### SARS-CoV-2 BA.4 and BA.5 exhibit different pathogenicity *in vivo*

With the implementation of vaccines, Omicron has become the dominant circulating SARS-CoV-2 strain (19). However, different Omicron subvariants exhibit unequal pathogenicity *in vivo* (17, 20–22). The viral S protein was identified to be among the major contributors to SARS-CoV-2 virulence (23). BA.4 and BA.5 are two closely related Omicron subvariants with similar S proteins. To examine whether BA.4 and BA.5 exhibit the same pathogenicity *in vivo*, K18-hACE2 mice were intranasally infected with natural BA.4 and BA.5 isolates. This experiment revealed that BA.4 and BA.5 had different virulence in K18-hACE2 mice. BA.4 was apathogenic, whereas BA.5 led to 100% lethality (**Figs. 1A and 1B**). Notably, while viral replication in the lungs was comparable between BA.5- and BA.4-infected animals (**Fig. 1C**), immunohistochemistry (IHC) analysis revealed higher levels of viral nucleocapsid (N) protein in BA.5-infected animals compared to BA.4 (**Fig. 1D**). Histopathological analysis showed increased cellular infiltrates in the lungs of BA.5-infected animals compared to BA.4-infected mice (**Fig. 1D**). Coinciding with these lung pathology results, we observed increased number of immune cells (**Supplemental Fig. 1A**) and cytokines (**Supplemental Fig. 1B**) in bronchoalveolar lavage fluids (BALFs) at 5 days post-infection (dpi). Specifically, monocyte derived macrophages, NK cells, γδT cells, NKT cells, CD4^+^ and CD8^+^ T cells were increased in BA.5-infected mice, whereas alveolar macrophages were significantly decreased compared to BA.4, possibly related to increased infection and induced cell death in this tissue resident cell population (**Supplemental Fig. 1A**). Likewise, proinflammatory cytokines IL-2, IL4, IL-6, IFN-γ, and TNF-α were significantly elevated in BALFs of BA.5-infected K18-hACE2 mice compared to those infected with BA.4 (**Supplemental Fig. 1B**). We found no significant pathological differences in the brains of animals infected with either BA.4 or BA.5, and no presence of viral N protein in the brains of animals infected with BA.4 or BA.5 (**Fig. 1E**). Altogehter, these results indicate that Omicron BA.5 infection caused more severe disease in K18-hACE2 mice than Omicron BA.4, suggesting that genetic differences between Omicron variants BA.5 and BA.4 may be responsible for the differences in viral pathogenicity.

**Figure 1.**
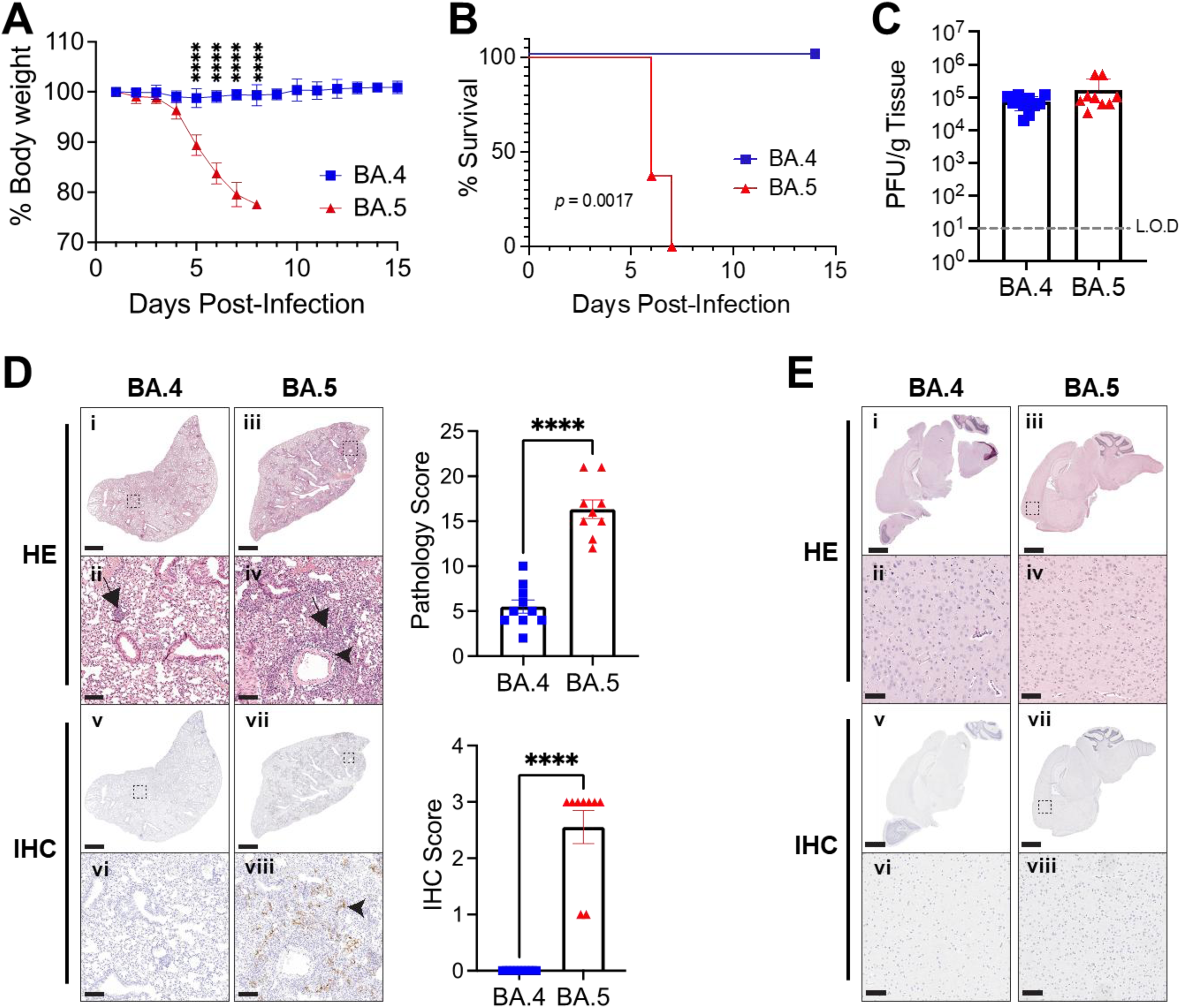
Omicron BA.4 infection displays a different outcome than Omicron BA.5 in K18- hACE2 mice: (**A**) **Body weight:** K18-hACE2 (5-8 months old) were infected intranasally (3.25×10^4^ PFU/mouse) with BA.4 (n=3, male) or BA.5 (n=8, female and male). Changes in body weight were determined daily for 14 days and plotted according to the ratio of the body weight each day over the body weight right before infection. Data are presented as Mean ± Standard Deviation (SD). Unpaired Student’s *t*-tests with Welch’s correction were used to compare the means between the indicated groups. ****, *p*<0.0001. (**B**) **Survival:** The survival curve was plotted according to the death of the mice in panel A. Animals losing more than 25% of their initial body weight were considered to reach the endpoint of the experiment. Survival differences between the groups of BA.4- and BA.5-infected mice were analyzed using the log-rank (Mantel-Cox) test. (**C**) **Viral titers:** Amount of virus in the lungs of K18-hACE2 mice infected intranasally (3.25×10^4^ PFU/mouse) with BA.4 (n=3) or BA.5 (n=8) at 5 dpi. L.O.D: limit of detection. (**D-E**) **HE (i-iv) and IHC (v-viii) analysis of the lungs (D) and brains (E) of mice infected in panel C.** Representative images of the lungs and brains of BA.4- or BA.5-infected K18-hACE2 mice at 5 dpi. IHC detection of viral N protein (arrowhead), along with interstitial (arrow) and perivascular inflammatory infiltrates (arrowheads), was more prominent in lung tissues of mice infected with BA.5 compared to those infected with BA.4. Scale bars: 1 mm (i, iii, v, vii); 100 μm (ii, iv, vi, viii). The pathology score and the IHC score of the lungs are indicated. Data are presented as Mean ± SD. Mann-Whitney tests were used to compare the differences between groups. ****, *p*<0.0001.

### Establishment of a reverse genetic system for BA.5

To address the different pathogenicity of Omicron BA.4 and BA.5, we established a bacterial artificial chromosome (BAC)-based reverse genetic system (RGS) for BA.5. To this end, we divided the BA.5 viral genome into 5 fragments that were chemically synthesized and assembled into an empty BAC plasmid using unique restriction sites and standard molecular biology approaches (**Fig. 2A**). After assembly of the entire BA.5 viral genome we analyzed the BAC by restriction enzyme and gel electrophoresis (**Fig. 2B**). Then, the recombinant (r)BA.5 was rescued by transfection of the BAC into Vero AT cells. We further characterized the rBA.5 in cultured Vero AT cells by assessing and comparing with a natural BA.5 isolate by plaque assay (**Fig. 2C**) and growth kinetics (**Fig. 2D**). Both rBA.5 and BA.5 formed uniform plaques of similar sizes and morphology (**Fig. 2C**). Likewise, the rBA.5 exhibited similar growth kinetics and peak titers than the natural BA.5 isolate (**Fig. 2D**). These results demonstrate that the rBA.5 generated using BAC-based RGS has similar fitness than a natural Omicron BA.5 isolate in Vero AT cells.

**Figure 2.**
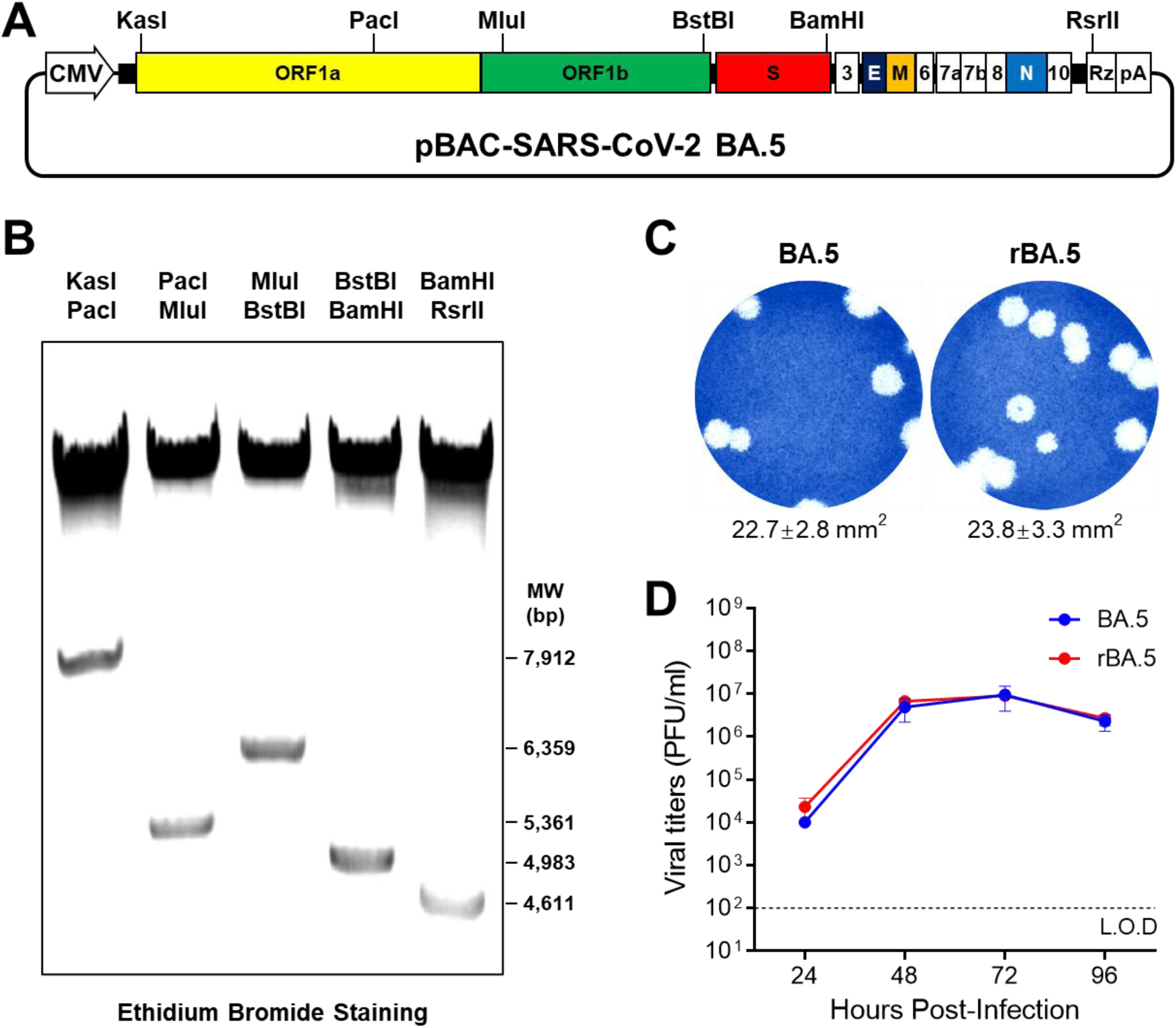
Establishment of a reverse genetic system for BA.5: (**A**) **Schematic diagram of the Omicron BA.5 BAC**: Unique restriction sites used to assemble the entire genome of Omicron BA.5 are indicated. Viral structural and accessory ORF proteins are shown. Not draw in scale. (**B**) **Restriction analysis of BA.5 BAC:** The BA.5 BAC was digested with the indicated restriction enzymes and separated on a 0.5% agarose gel. The image was photographed using a ChemiDoc imager. (**C**) **Plaque morphology:** Monolayers of Vero AT cells (6-well plate format) were infected with ∼10 PFU of BA.5 natural isolage (left) or the recombinant BA.5(right) for 1 h. After viral infection, cell monolayers were overlayed with semi-solidified media containing 0.5% agar and cultured at 37 °C for 72 h. Viral plaques were visualized with crystal violet staining. (**D**) **Growth kinetics:** Monolayers of Vero AT cells (6-well plate format, triplicates) were infected (MOI 0.001) with BA.5 or recombinant BA.5 for 1h, and cell culture supernatants (CCS) were collected at the indicated time points. Viral titers were plotted according to the titers in the CCS determined by plaque assay. Data are presented as Mean ± Standard Error of Mean (SEM).

### *In vivo* characterization of rBA.5

Next, we compared the *in vivo* viral dynamics and pathogenicity of rBA.5 with a BA.5 natural isolate in K18-hACE2 mice. Mice were intranasally mock-infected with PBS or infected with 3.75×10^5^ PFU/mouse of BA.5 or rBA.5. Consistent with our previous observations (17) (**Fig. 1**). BA.5 infected mice started losing weight at 2 dpi with significant differences observed at 3-6 dpi compared to PBS mock-infected controls (**Fig. 3A**). Similarly, rBA.5 infected mice followed the same weight loss trail and showed significant weight loss at 3-6 dpi (**Fig. 3A**). One mouse of each infection group recovered and survived the infection with similar recovery weight kinetics (**Fig. 3A**). Mortality was observed in both rBA.5 and BA.5 infected mice starting at 5 dpi, with 87.5% of the rBA.5 and BA.5 infected mice succumbing to infection by 5 or 6 dpi (**Fig. 3B**). Subsequently, we compared lung viral titers from the mice that succumbed to infection at 6 dpi. Similar levels of infectious virus titers were observed in the lungs of both rBA.5 and BA.5 infected mice (**Fig. 3C**). This was confirmed by immunolabeling against the SARS-CoV-2 N protein (**Fig. 3D**). Histopathological analysis of the lungs at 6 dpi in both rBA.5 and BA.5 infected mice displayed hyperplasia of type II pneumocytes, increased interstitial and perivascular infiltrates, alveolar cells, and edema (**Fig. 3E**), demonstrating that, *in vivo* viral dynamics and pathology were comparable between BA.5 *vs.* rBA.5 in K18-hACE2 infected mice.

**Figure 3.**
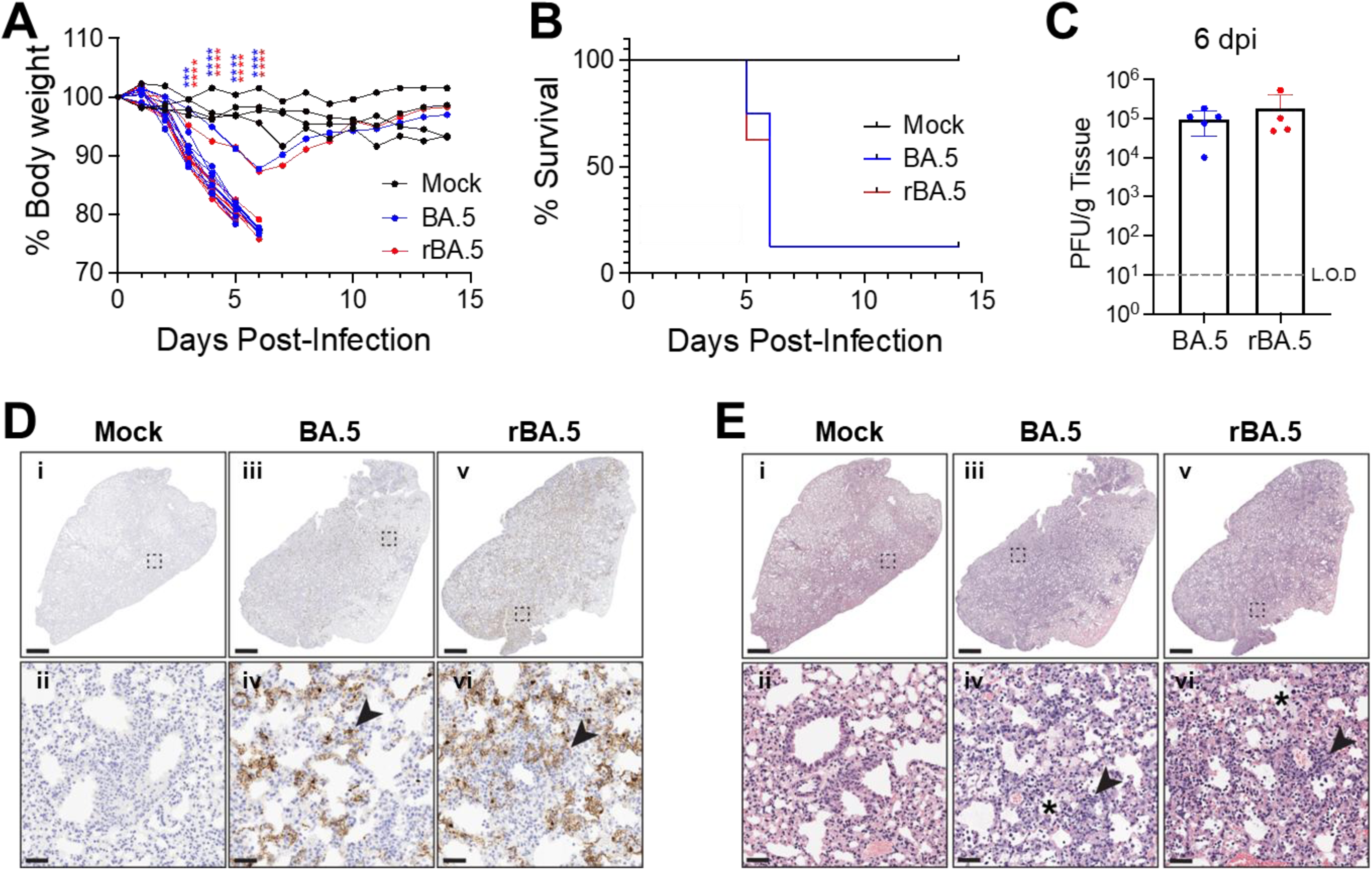
rBA.5 has similar pathogenicity than BA.5 in K18-hACE2 mice: (**A**) ) **Body weight:** K18-hACE2 mice (5-8 months old) were infected intranasally (3.75×10^5^ PFU/mouse) with BA.5 (n=8: 4 male, 4 female) or rBA.5 (n=8: 4 male, 4 female), or equivalent volume of PBS (Mock, n=4: 2 male, 2 female). Changes in body weight were monitored daily for 14 days and plotted according to the ratio of the body weight each day over the body weight right before infection. Two-way ANOVA with Tukey’s multiple comparisons test was used to compare the differences among groups. Data are represented as the mean ± SD. ***, *p*< 0.001; ****, *p*< 0.0001. (**B**) **Survival:** Percent survival of K18- hACE2 mice following infection with BA.5 (n=8: 4 male, 4 female), rBA.5 (n=8: 4 male, 4 female), or PBS (Mock, n=4: 2 male, 2 female). Animals losing more than 25% of their initial body weight were considered to reach the endpoint of the experiment. The log-rank (Mantel-Cox) test was used to compare the difference between groups of BA.5 and rBA.5. (**C**) **Viral titers:** Viral titers in the lungs of the K18-hACE2 mice infected with BA.5 (n=8) or rBA.5 (n=8) were assessed at 6 dpi. L.O.D: limit of detection. (**D**) Representative IHC staining (i-vi) targeting the viral nucleoprotein (arrowheads) shows similar antigen distribution in both groups at 6 dpi. Scale bars: 1 mm (i, iii, v); 50 μm (ii, iv, vi). (**E**) Perivascular (arrowheads) and alveolar (asterisks) infiltrates of representative HE (i-vi) stained lung sections were comparable between BA.5 and rBA.5 groups at 6 dpi. Scale bars: 1 mm (i, iii, v); 50 μm (ii, iv, vi).

### KSF amino acid deletion attenuates NSP1 host translation shutoff

To better understand potential drivers for the differences in pathogenicity between variants BA.4 and BA.5, a complete amino acid comparison was performed. Several differences were found. Notably, one of the few differences between BA.4 and BA.5 consists in triple deletion of the amino acids lysine, serine and phenylalanine at 141-143, hereafter depicted as ΔKSF, in BA.4 NSP1 (**Fig. 4A**). NSP1 contains a N-terminal domain (NTD), a linker domain, and a C-terminal domain (CTD) (**Fig. 4B**), and both NTD and CTD were reported to be involved in inhibition of host gene translation by different mechanisms (24). Importantly, the linker domain in NSP1 of SARS-CoV and the ancestral SARS-CoV-2 is conserved (**Fig. 4C**). To evaluate the ability of BA.4 and BA.5 NSP1 proteins to inhibit host gene expression, we used a previously described plasmid-based shutoff assay (25). We found that both BA.4 and BA.5 NSP1 inhibit expression of EGFP and Gaussia luciferase (Gluc) in a dose-dependent manner; however, BA.4 NSP1 inhibition of host gene translation was significantly lower than that of BA.5 NSP1 (**Figs. 4D and 4E**). Collectively, these data demonstrate that the ΔKSF in BA.4 NSP1 results in attenuated host translational shutdown compared to BA.5 NSP1.

**Figure 4.**
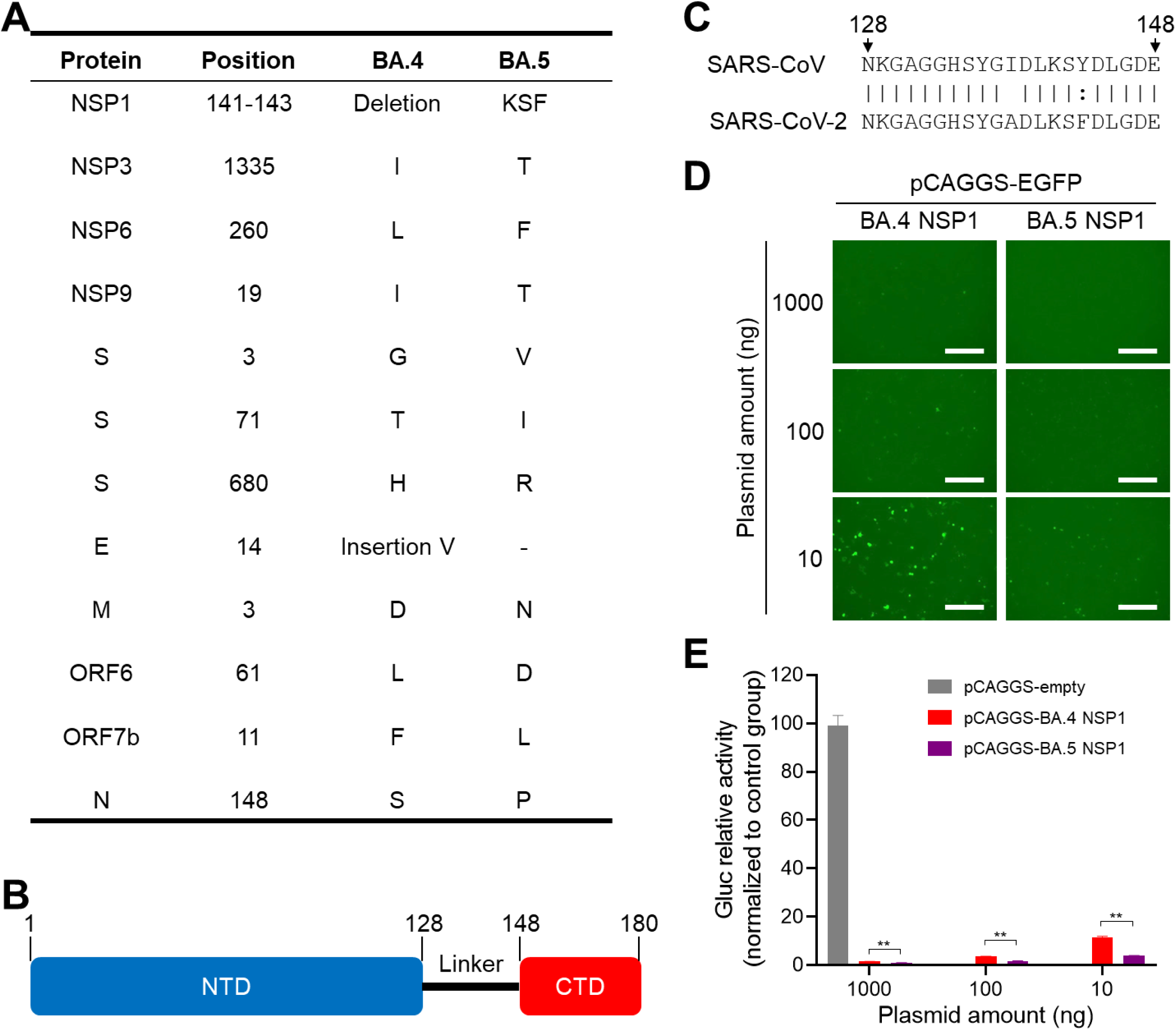
Inhibition of host gene expression by BA.4 and BA.5 NSP1: (**A**) **Genomic comparison of BA.4 and BA.5**: Amino acid differences in the viral proteins between BA.4 and BA.5 are indicated. Amio acid positions were referenced to the original SARS-CoV-2 WA1 strain. (**B**) Schematic representation of NSP1 domains: N-terminal domain (NTD), linker, and C-terminal domain (CTD) of NSP1 are represented, including their amino acid length. (**C**) **Alignment of NSP1 linker domains of SARS-CoV and SARS-CoV-2:** Double dots indicate high similarity. Bars show identical amino acids. Empty spaces indicate different amino acids. (**D-E**) **Inhibition of host gene expression:** Monolayers of HEK 293T cells (12-well format, triplicates) were co-transfected with pCAGGS-EGFP (500 ng/well) and pCAGGS-Gluc (500 ng/well), along with the indicated amounts of pCAGGS NS! plasmids for BA.4 and BA.5. To maintain a consistent total plasmid amount (2,000 ng/well) across all wells, pCAGGS empty plasmid was used. At 48 h post-transfection, EGFP expression was assessed under a fluorescent microscope, and representative images are shown (**D**). Meanwhile, 25 µl of CCS were collected, and Gluc activity was measured using a luciferase plate reader (**E**). Scale bars: 100 µm. Data are presented as Mean ± SD. Unpaired Student’s *t*-tests with Welch’s correction were used to compare the means between the indicated groups. **, *p*<0.01.

### NSP1ΔKSF attenuates BA.5 pathogenicity *in vivo*

To test whether ΔKSF in NSP1 contributes to the difference in pathogenicity between BA.4 and BA.5, we introduced ΔKSF into the BA.5 genome and engineered a rBA.5 NSP1ΔKSF loss-of-function mutant. By comparing the characteristics in cultured Vero AT cells, rBA.5 NSP1ΔKSF formed smaller-sized plaques (**Fig. 5A**) but exhibited similar growth kinetics to rBA.5 (**Fig. 5B**). Next, we compared the *in vivo* characteristics of rBA.5 and rBA.5 NSP1ΔKSF in K18-hACE2 mice. Mice were intranasally infected with rBA.5 or rBA.5 NSP1ΔKSF and body weights were monitored daily. Notably, we observed a significant difference in changes in body weight in mice infected with rBA.5 and rBA.5 NSP1ΔKSF starting at 3 dpi up to 14 dpi, with rBA.5 infected mice succumbing to infection starting at 5 dpi, whereas the rBA.5 NSP1ΔKSF infected mice started to lose weight at 6 and 7 dpi and had almost fully recovered by 14 dpi (**Supplemental Fig. 2A**). Overall, 100% of the rBA.5 infected mice succumbed to infection by 6 dpi whereas only 10% of the rBA.5 NSP1ΔKSF infected mice succumbed to infection (**Supplemental Fig. 2B**). Viral load assessment in the lungs upon death or at the study endpoint (14 dpi) showed the presence of the infectious virus at humane endpoint and complete viral clearance in surviving in rBA.5 NSP1ΔKSF infected mice (**Supplemental Fig. 2C**). Consistent with the viral titer result, IHC for the viral N protein confirmed the presence of viral antigen in rBA.5 infected animals at humane endpoint and no viral antigen in the rBA.5 NSP1ΔKSF infected animals (**Supplemental Figs. 2D and 2F**). Histopathological analysis of lung tissue revealed severe pathology in the rBA.5 group characterized by increased perivascular and interstitial inflammation, type II pneumocytes, alveolar cell hyperplasia, alveolar edema and bronchiolar cell alterations at humane endpoint; in contrast, animals infected with rBA.5 NSP1ΔKSF displayed low levels of pathology at study endpoint (**Supplemental Figs. 2E and 2F, and Supplementary Table 1**). Thus, the NSP1ΔKSF mutation significantly attenuates BA.5, leading to delayed body weight loss and increased survival in K18-hACE2 mice.

**Figure 5.**
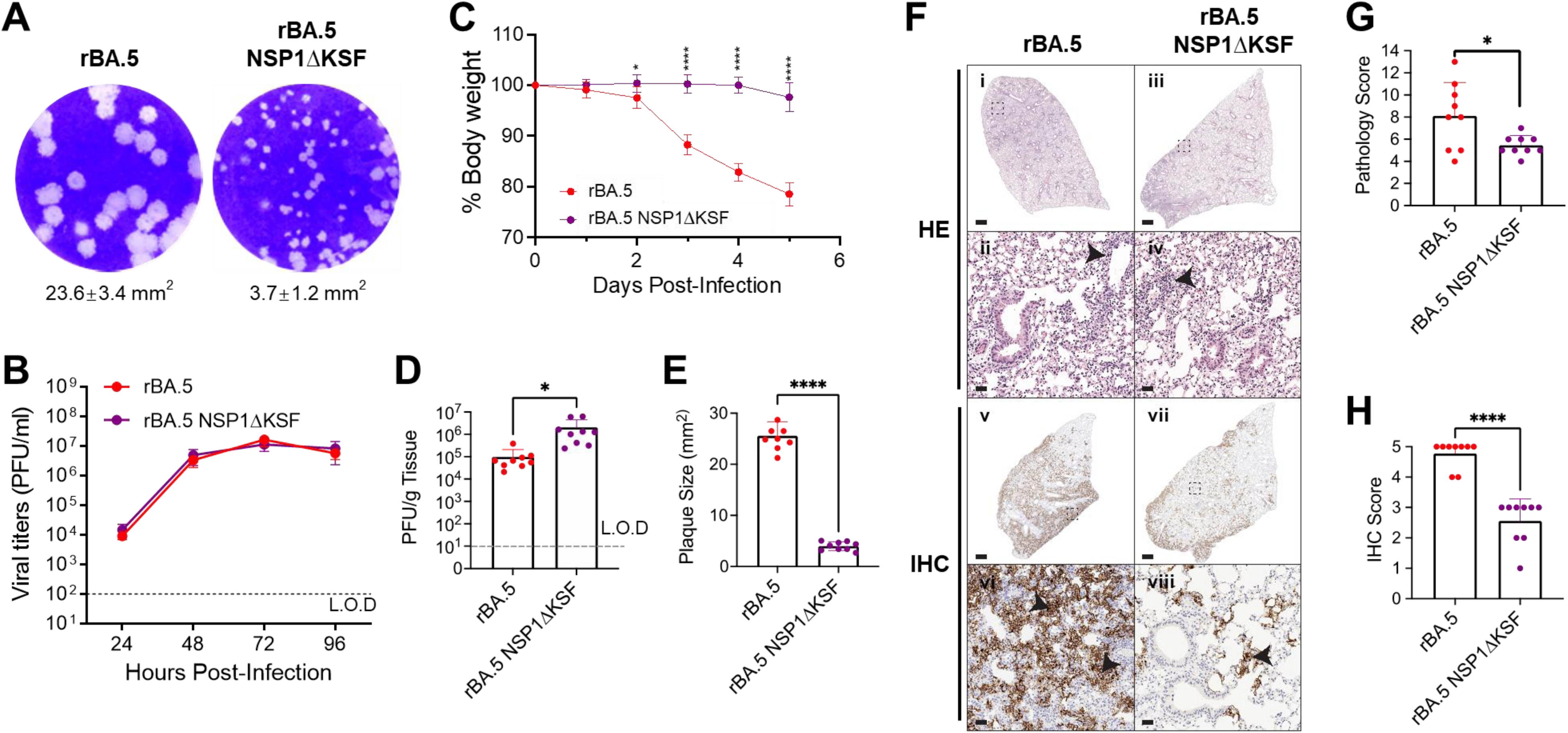
KSF deletion attenuates BA.5 pathogenicity *in vivo*: (**A**) **Plaque morphology**: Plaque assay was performed by infecting monolayers of Vero AT cells (6-well plate format) with rBA.5 (left) or rBA.5 NSP1ΔKSF (right) as previously described. Viral plaques were stained with crystal violet, and plaque sizes were measured using ImageJ. (**B**) **Growth kinetics:** Vero AT cell monolayers (6- well plate format, triplicates) were infected with rBA.5 or rBA.5 NSP1ΔKSF at an MOI of 0.001 for 1 h. After viral infection, the supernatant was replaced with post-infection media (3 mL/well). The CCS were then collected at the indicated time points, and viral titers were determined by plaque assay. Data are presented as Mean ± SEM. (**C-H**) ***In vivo* studies:** K18-hACE2 mice (5-8 months old) were infected with rBA.5 or rBA.5 NSP1ΔKSF (n=9: 5 male, 4 female per group) and monitored for body weight changes (**C**). Student’s *t*-test was used to compare the means between the indicated groups. *, *p*<0.05; ****, *p*<0.0001. Mice were sacrificed at 5 dpi, and lungs were collected to assess viral titers (**D**). L.O.D: limit of detection. Data are presented as Mean ± SEM. An unpaired Student’s *t*-test with Welch’s correction was used to compare the means between the indicated groups. *, *p*<0.05. Plaque size from lung homogenates of rBA.5 and rBA.5NSP1ΔKSF infected mice at 5 dpi was measured using ImageJ (**E**). Data are presented as Mean ± SD. An unpaired Student’s *t*-test was used to compare the means between the indicated groups. ****, *p*<0.0001. HE (i-iv) and IHC for nucleoprotein (v-viii) representative images of lungs collected at 5 dpi are shown (**F**). HE-stained and IHC sections show lung pathology and viral antigen distribution in rBA.5- and rBA.5 NSP1ΔKSF- infected mice. Perivascular inflammatory infiltrates (arrowheads in ii and iv), along with IHC staining for the viral nucleoprotein (arrowheads in vi and viii), appeared more pronounced in lungs from rBA.5-infected mice. Scale bars: 1 mm (i, iii, v, vii); 100 μm (ii, iv, vi, viii). Pathology scoring (**G**) and IHC scoring (**H**) were performed on the lung sections at 5 dpi. Data are presented as Mean ± SD. Mann-Whitney tests were used to compare differences between groups for the scores of pathology and IHC. *, *p*<0.05; ****, *p*<0.0001.

Upon confirming that NSP1 KSF deletion attenuates the pathogenicity of the rBA.5 and increases survival rates, we compared the viral titers and pathology in the lungs for all animals at 5 dpi to better understand the potential factors driving the differential outcomes. K18-hACE2 mice were intranasally inoculated with either rBA.5 or rBA.5 NSP1ΔKSF. At 5 dpi, rBA.5 infected mice displayed an average of 20% body weight loss, while those infected with rBA.5 NSP1ΔKSF showed only a 3% decrease (**Fig. 5C**). Lung viral titers were one-log higher in rBA.5 NSP1ΔKSF infected mice as compare to rBA.5 infected mice (**Fig. 5D**). However, plaque sizes, which positively correlate with increased virulence (17, 23), were 6.4 times smaller for rBA.5 NSP1ΔKSF infected mice compared to rBA.5 infected mice (**Fig. 5E**). Immunolabeling for the N protein showed a 45% reduction in viral antigen in the rBA.5 NSP1ΔKSF group compared to the rBA.5 group (**Figs. 5F and 5G**). Lung histopathology showed mild interstitial inflammation in both groups, but mice infected with rBA.5 revealed more severe histopathological changes characterized by increased perivascular inflammation and type II pneumocyte and alveolar cell hyperplasia (**Fig. 5F and Supplementary Table 2**). On average, rBA.5 NSP1ΔKSF infected mice had a 31% lower pathology score than the rBA.5 group (**Fig. 5H**). Together, these data suggest that the NSP1ΔKSF mutation significantly reduces the pathogenicity of BA.5 in K18-hACE2 mice, indicating a substantial reduction in virulence.

### KSF deletion weakens NSP1-mediated inhibition of host gene expression and reduces WA1 pathogenicity *in vivo*

To further confirm the contribution of KSF deletion in the inhibition of host gene expression, we introduced the KSF deletion into the original WA1 strain NSP1. Using the plasmid-based shutoff assay, we found that both WA1 NSP1 and WA1 NSP1ΔKSF inhibited EGFP and Gluc expression in a dose-dependent manner; however, WA1 NSP1ΔKSF exhibited impaired inhibition of host gene expression compared to WA1 NSP1 (**Figs. 6A, 6B, and 6C**). To demonstrate the contribution of NSP1 KSF deletion to viral virulence, we introduced this deletion into the WA1 genome and rescued a rWA1 NSP1ΔKSF. The rWA1 NSP1ΔKSF formed plaques of similar size (**Fig. 7A**) and exhibited comparable growth kinetics (**Fig. 7B**) to rWA1 in Vero AT cells. K18-hACE2 mice infected with rWA1 NSP1ΔKSF and rWA1 began losing weight at 2 dpi (**Fig. 7C**). However, while all mice infected with rWA1 succumbed to infection by 7 dpi, only one out of five mice succumbed to rWA1 NSP1ΔKSF infection (**Fig. 7D**). Both viruses displayed similar replication in the lungs at 2 and 4 dpi (**Fig. 7E**), but rWA1 NSP1ΔKSF exhibited slightly slower replication in the nasal turbinates at both times post- infection (**Fig. 7F**). These findings confirm our previous observation with BA.5 and demonstrate that KSF deletion in WA1 NSP1 also results in viral attenuation without significantly impacting viral replication *in vitro*.

**Figure 6.**
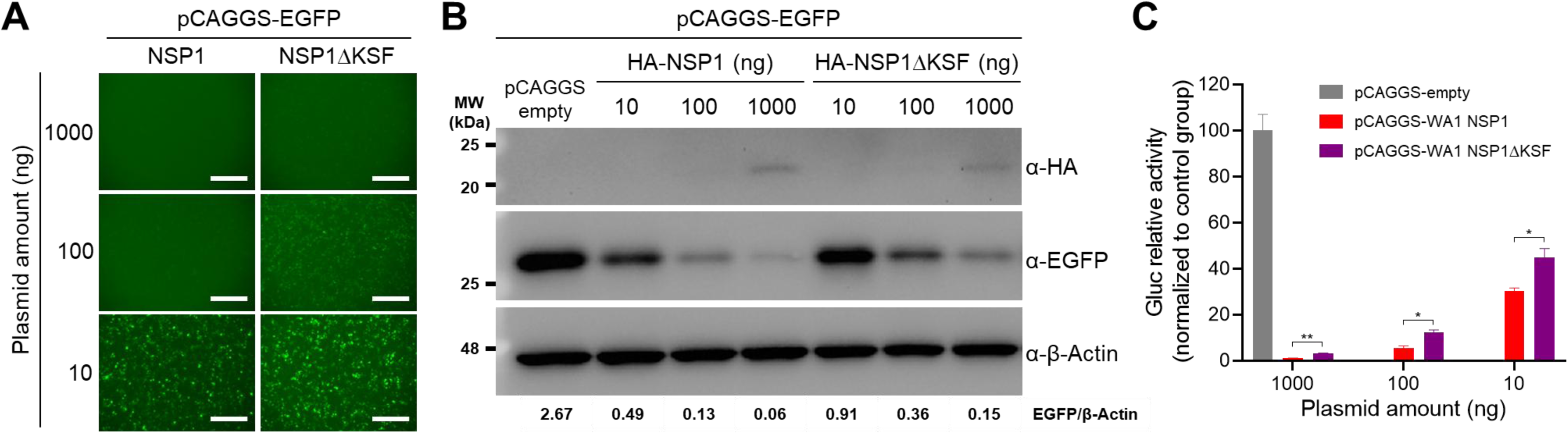
KSF deletion attenuates WA1 NSP1 inhibition of host gene expression: HEK 293T cell monolayers (12-well format, triplicates) were co-transfected with pCAGGS-EGFP (500 ng/well) and pCAGGS-Gluc (500 ng/well), along with the indicated amounts of pCAGGS WA1 NSP1 WT or NSP1ΔKSF expression plasmids. To ensure a consistent total plasmid amount (2,000 ng/well) across all wells, pCAGGS empty plasmid was used. At 48 h post-transfection, EGFP expression was assessed under a fluorescent microscope, with representative images shown (**A**). Scale bars: 100 µm. Cell lysates from transfected cells were used to evaluate NSP1, EGFP, and β-Actin expression by Western blot (**B**). The relative EGFP expression normalized to β-Actin was quantified using ImageJ. Meanwhile, 25 µL of CCS were collected from each of the wells and Gluc activity was measured using a luciferase plate reader (**C**). Data are presented as Mean ± SD. Unpaired Student’s *t*-tests with Welch’s correction were used to compare the means between the indicated groups. *, *p*<0.05; **, *p*<0.01.

**Figure 7.**
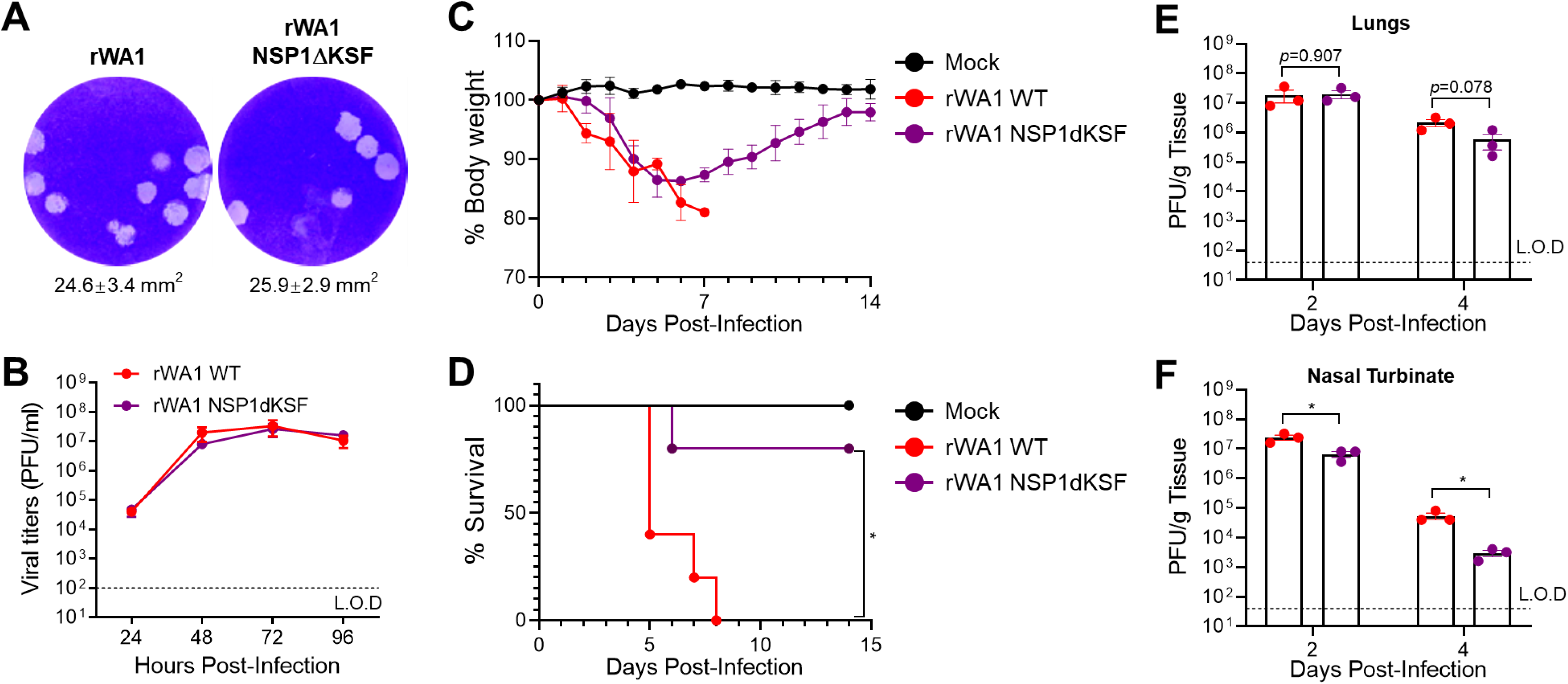
KSF deletion in NSP1 attenuates SARS-CoV-2 WA1 *in vivo*: (**A**) **Plaque morphology:** Plaque assays wereperformed by infecting monolayers of Vero AT cells (6-well plate format) with rWA1 (left) or rWA1 NSP1ΔKSF (right). Viral plaques were visualized using crystal violet staining, and plaque size was measured using ImageJ. (**B**) **Growth kinetics:** Vero AT cell monolayers (6-well plate format, triplicates) were infected with rWA1 or rWA1 NSP1ΔKSF at an MOI of 0.0001 for 1 h. After viral infection, the supernatant was replaced by post-infection media (3 mL/well). CCS were collected at the indicated time points, and viral titers were determined by plaque assay. Data are presented as Mean ± SEM. (**C-D**) **Body weight and survival:** Eight-week-old female K18-hACE2 mice (n=5) were mock-infected with PBS or infected intranasally with 10^5^ PFU/mouse of rWA1 or rWA1 NSP1ΔKSF. Changes in body weight (**C**) and survival (**D**) were monitored daily until 14 dpi. Survival differences between the indicated groups were analyzed using the log-rank (Mantel-Cox) test. *, *p*<0.05. (**E-F**) **Viral titers**: Eight-week-old female K18-hACE2 mice (n=6) were infected intranasally with 10^5^ PFU/mouse of rWA1 or rWA1 NSP1ΔKSF. At 2 and 4 dpi, three mice per group were randomly selected, and lungs and nasal turbinate were harvested. Viral titers in the lungs (**E**) and nasal turbinate (**F**) were determined by plaque assay. L.O.D: limit of detection. Data are presented as Mean ± SEM. Statistical significance was determined using Student’s *t*-tests between the indicated groups. *, *p*<0.05.

## DISCUSSION

Here, we provide data demonstrating that Omicron BA.5 is significantly more pathogenic than the closely related Omicron BA.4 in K18-hACE2 mice. Despite comparable levels of infectious virus between Omicron BA.4- and BA.5-infected mice at 5 dpi, IHC analysis revealed an increased presence of viral N protein in the lungs of BA.5-infected mice, suggesting a greater accumulation of viral antigen in these tissues. These findings suggest that Omicron BA.5 established a more robust infection than Omicron BA.4, with BA.5 causing disease and mortality in K18-hACE2 mice, whereas BA.4 resulted in attenuated disease and no mortality. However, neither Omicron BA.5 nor BA.4 led to neurological symptoms, similar to what we and other groups have reported before (17, 26). The process of viral propagation can result in mutations that either enhance or diminish pathogenesis

(27, 28). It is plausible that the severe disease and pronounced pathology associated with Omicron BA.5 stem from its ability to establish infection more rapidly than BA.4, potentially triggering immune hyperactivation that can lead to cytokine release syndrome (CRS) and acute respiratory distress syndrome (ARDS), both of which are linked to severe outcomes in SARS-CoV-2 infections (29). To gain deeper insights into the role of the immune response in the observed pathology, we analyzed the immune profiles of animals infected with Omicron BA.4 and BA.5 at 5 dpi. Inflammatory cells, including neutrophils, monocyte-derived macrophages, natural killer (NK) cells, and activated T cells, were notably elevated in the BALFs of BA.5-infected animals. Although these cells play a crucial role in viral clearance, their overactivation has been implicated in contributing to disease severity and pathology (30–33). Moreover, pro-inflammatory cytokines such as IL-6, TNF-α, IL-2, IFN-γ, and IL-4 were found to be elevated in the BALFs. While these cytokines likely play a role in initiating an antiviral immune response in BA.5-infected mice, they may simultaneously exacerbate inflammation and contribute to CRS. The increased pathology observed in BA.5 compared to BA.4 infection can likely be due to increased immune cell infiltrates and proinflammatory cytokines seen in the BALFs of these animals (34). However, we cannot conclude from these data alone whether the inflammation and pathology are linked to the increased virulence of BA.5 or whether other immunosuppressive mechanisms were at play.

Omicron BA.5 has acquired several mutations that may contribute to the observed differences in pathogenicity compared to Omicron BA.4. Our analysis revealed 12 mutations in BA.5 that distinguish it from BA.4, including three in the spike (S), one in the envelope (E), one in the membrane (M), and another one in the N structural proteins; one in the ORF6, and another one in the ORF7b; and the remaining changes in NSP1, NSP3, NSP6, and NSP9. The S, which is essential for viral entry into host cells, plays a critical role in infection but may have only limited contributions to overall pathogenesis, particularly in the acute stages of disease (35). Other viral proteins, however, have been shown to influence disease outcomes significantly (36, 37). Several SARS-CoV-2 proteins are involved in modulating the immune response, including NSP1, NSP3, NSP6, ORF6, and the M protein, which collectively inhibit IFNβ (38–40). Considering the more severe lung pathology and clinical outcomes associated with BA.5, it is plausible that changes in NSP1 enhance its ability to shut down host cell translation as compared with BA.4, leading to greater tissue damage. This heightened damage may activate the immune system, resulting in CRS, exacerbated pathology, and, ultimately, more severe disease outcomes.

Given the differences in pathogenicity between BA.4 and BA.5, we developed a reverse genetics system to investigate the underlying drivers of Omicron BA.5’s increased pathogenicity. To avoid gain-of-function concerns, we conducted a loss-of-function approach to understand the pathogenicity discrepancy between BA.4 and BA.5. We focused on the effects of NSP1 on the inhibition of host cell translation, and demonstrated that the KSF deletion in NSP1 is a major contributor to BA.5 virulence. All viruses rely on the host cellular ribosomes to translate their mRNAs into polypeptides. A common viral strategy to achieve this is to impair the translation of cellular mRNAs in a process termed “host translational shutoff”. Host cell translational shutoff redirects translational resources, specifically ribosomes, towards viral RNAs, thereby blocking the host cell from mounting an efficient antiviral response and aiding assembly of new progeny virons (41). Several SARS-CoV-2 proteins have previously been implicated in shutting off host translation. For instance, ORF6 imprisons the host mRNA in the nucleus by clogging the nuclear pore through its interactions with the mRNA export factors, Rae1 and Nup98 (42, 43). NSP8 and NSP9 bind to the 7SL RNA in the signal recognition particle and interfere with protein trafficking to the cell membrane upon infection, and NSP16 binds to the mRNA recognition domains of the U1 and U2 splicing RNAs and disrupts mRNA splicing (42, 44). However, one of the first proteins produced during SARS-CoV- 2 infection, NSP1, is considered the major contributor to host cell shutoff (45). There are three elucidated mechanisms by which NSP1 inhibits host gene translation: (1) inhibition of the 40S ribosome by binding of the NSP1 CTD to the 40S mRNA entry site to block the mRNA entry tunnel through multiple side chains such as Y154, F157, W161, K164, H165, R171, and R175 (40, 46); (2) binding to 18S rRNA in the mRNA entry channel of the 40S ribosome (44), (3) interaction with the host mRNA export receptor heterodimer NXF1-NXT1 to prevent proper binding of NXF1 to mRNA export adaptors and NXF1 docking at the nuclear pore complex (47); and (4) by stabilizing the NSP1 NTD binding to the 40S ribosomal subunit to promote mRNA decay (48–50). In this study, we found that, beyond the previously studied roles of NSP1 CTD and NTD in inhibition of host gene expression, three amino acids (KSF) within the linker domain also contribute to NSP1-mediated host cell translation shutoff. However, structural studies mapping how the KSF deletion impacts NSP1 interaction with the 40s ribosome or heterodimer NXF1-NXT1 are needed to better understand the underlying mechanism of host inhibition.

In addition to the host cell translation shutoff observed *in vitro*, *in vivo* studies demonstrated reduced virulence by both WA-1 and BA.5 expressing NSP1ΔKSF. We observed a substantial difference in infection outcomes between rBA.5 and rBA.5 NSP1ΔKSF, underscoring the role of the KSF deletion in NSP1 in modulating viral protein translation *in vivo*. This finding was corroborated by reduced viral antigen staining in lung tissues in mice infected with rBA.5 NSP1ΔKSF. Interestingly, despite the attenuation of clinical symptoms in these mice, viral titers in the lungs were higher at 5 dpi compared to rBA.5-infected mice. However, rBA.5 NSP1ΔKSF-infected cells produced smaller plaques, a phenotype associated with reduced virulence (51). The KSF deletion likely alters the structural conformation of the NSP1 linker domain, modulating its functional interaction with the host translation machinery. This modification appears to partially affect NSP1’s ability to suppress host protein synthesis, thereby restoring a balance between host and viral protein translation. As a result, viral protein synthesis may be deprioritized, enabling the translation of critical host proteins involved in cellular homeostasis and immune responses. This rebalancing could account for the reduced cytopathic effect observed *in vivo*, potentially mitigating the severity of inflammatory responses and immunopathology. This is consistent with prior research demonstrating that SARS-CoV-2 pathogenesis is closely linked to the interplay between viral protein synthesis and host immune activation.

Additionally, we observed increased pathology in the BA.5 and rBA.5 infected mice, which is consistent with previous observations in which SARS-CoV-2 infection can induce immune hyperactivation, resulting in increased pathology and mortality (17, 34). Our present and previous results demonstrate a role of NSP1 and ORF6 mutations present in BA.4 in modulating severe disease with a minimal impact on virus replication *in vivo*, which may explain their ability to transmit in humans (18). Future studies will be needed to investigate the roles of other gene differences observed between BA.4 and BA.5, and the underlying mechanisms by which they may drive the differences observed alone or in combination with NSP1.

The KSF deletion found in BA.4 NSP1 but not in BA.5 and even in the latest XBB.2.3.3 variant indicates that the emergence of the KSF deletion in BA.4 NSP1 may not be positively selected under the pressure of pre-existing immunity due to previous infections or vaccinations. Meanwhile, because of its contribution to viral pathogenesis, NSP1 represents a reasonable target for the design of attenuated forms of SARS-CoV-2, alone or in combination with other attenuation markers, for their rational design of live-attenuated vaccines. Previously, we demonstrated that NSP1 mutations that impact its ability to inhibit the nuclear-cytoplasmic export of host mRNAs reduce the virulence of SARS-CoV-2 (52). In the current study, we demonstrate that mutations in NSP1 are associated with its second function, i.e., inhibition of mRNA translation, which also decreases virulence. Likewise, small molecule drugs that target the KSF region of NSP1 represent a reasonable target for the development of therapeutics against SARS-CoV-2.

## MATERIALS AND METHODS

### Biosafety

All the experiments using SARS-CoV-2 were approved by the Institutional Biosafety (IBC) and Institutional Animal Care and Use (IACUC) committee at Texas Biomedical Research Institute and Cornell University. All the *in vitro* and *in vivo* experiments with infectious natural isolates and recombinant SARS-CoV-2 were conducted at biosafety level 3 (BSL3) and animal BSL3 (ABSL3) laboratories, respectively, at Texas Biomedical Research Institute or Cornell University.

### Cells and viruses

Vero AT cells (Vero E6 cells expressing ACE2 and TMPRSS2) were obtained from BEI Resources (NR-54970) and maintained in DMEM supplemented with 10% FBS (Fetal Bovine Serum, VWR), 1% PSG (Penicillin-Streptomycin-Glutamine, Corning), and 10 µg/ml puromycin (Thermo Fisher Scientific). HEK 293T cells were maintained in DMEM supplemented with 10% FBS and 1% PSG. Natural isolates BA.4 (NR-56806) and BA.5 (NR-58620) were obtained from BEI Resources and expanded in Vero AT cells. Natural and recombinant SARS-CoV-2 were deep sequenced and sequences were confirmed before use.

### Recombinant viruses

The BA.5 genome was divided into 5 fragments, chemically synthesized (BioBasic), and assembled using unique restriction sites into an empty BAC plasmid as previously described (53). After restriction analysis verification, the BAC containing the full-length BA.5 genome was confirmed by sequencing (Plasmidsaurus). The KSF deletion in BA.5 NSP1 was introduced into a shuttle plasmid, and then the fragment containing KSF deletion in BA.5 NSP1 was released from the shuttle plasmid and used to replace the fragment between KasI and PacI in the BA.5 BAC. The resultant BAC was confirmed by sequencing (Plasmidsaurus). Similarly, the KSF deletion in NSP1 was also introduced into WA1 backbone BAC. Recombinant viruses were rescued by transfecting the corresponding BAC into Vero AT cells according to the previous protocol (54).

### Next-generation sequencing (NGS)

Trizol LS reagent (Thermo Fisher Scientific) was used to inactivate and extract RNA from the cell culture supernatant (CCS) of SARS-CoV-2 infected samples. The complete genome sequences of Omicron BA.4 and BA.5 viral stocks were obtained using a tailed amplicon approach as outlined in prior studies (17, 55). After performing multiplex PCR, libraries were prepared using the Twist Library Preparation EF Kit 2.0 alongside the Twist CD Index Adapter set 1-96 (Twist Bioscience). Sequencing was conducted on the Illumina iSeq platform with an iSeq 100 flow cell and cartridge. Raw sequencing data were processed using Trimmomatic for read trimming (56) and subsequently aligned to the SARS-CoV-2 reference genome (accession number OP295757) using the BWA- MEM2 tool (57) within the Galaxy platform (58).

### Amino acid comparison between BA.4 and BA.5 proteins

Consensus genomic sequences of Omicron BA.4 and BA.5 were obtained from NGS data. Amino acid positions were referenced against the WA1 strain (GenBank: MN985325.1). The comparison of viral genes between BA.4 and BA.5 was performed using the Basic Local Alignment Search Tool (BLAST).

### Plaque assays

Plaque assays were performed as previously described (59). Briefly, confluent monolayers of Vero AT cells (6-well plate format) were infected with natural isolate or recombinant SARS-CoV-2 at 37°C for 1 h. After viral adsorption, virus inoculum was discarded, cells were washed 3 times with PBS, overlaid with a semi-solid medium containing 0.5% agar, and incubated at 37°C for 72 h. After 24 h of fixation with 10% formalin, plates were taken from the BSL3 laboratory and stained with crystal violet. Plaque forming units (PFU) were determined based on plaque counts, and plaque sizes were measured using ImageJ (Fuji).

### Growth kinetics

Confluent monolayers of Vero AT cells (6-well plate format, triplicates) were infected (MOI 0.001) with the indicated SARS-CoV-2. After 1 h of virus adsorption at 37°C, cells were washed 3 times with PBS and incubated in post-infection media (DMEM containing 2.5% FBS and 1% PSG) at 37°C. At the indicated times post-infection, CCS were collected, and viral titers were determined by plaque assay.

### Western blot

Whole cell lysates, sodium dodecyl sulfate polyacrylamide gel electrophoresis (SDS-PAGE), and Western blot were performed as previously described (60). Briefly, cells were lysed in passive lysis buffer (Promega) at 4°C for 30 min, followed by centrifugation at 12,000 × *g* at 4°C for another 30 min. Equivalent amounts of cell lysates were subjected to 12% SDS-PAGE and transferred to nitrocellulose membranes. After blocking with 5% bovine serum albumin in PBS containing 0.1% Tween 20 at room temperature for 1 h, the membranes were incubated with primary antibodies at 4°C overnight, followed by horseradish peroxidase-conjugated secondary antibody incubation at 37°C for 1 h. An antibody against β-Actin was used as the loading control. Membranes were developed with ECL detection reagents (Thermo Fisher Scientific) in the ChemiDoc MP Imaging System (Bio-Rad).

### NSP1 host shutoff assay

Plasmids pCAGGS-EGFP (500 ng/well) and pCAGGS-Gluc (500 ng/well) were co-transfected with the indicated amounts of pCAGSS-NSP1 into HEK 293T cells (12-well format, triplicates). The amount of total plasmids was kept consistent (2,000 ng/well) by using a pCAGGS empty plasmid. At 48 h post-transfection, GFP expression was determined under an inverted fluorescent microscope, and Gluc activity in CCS was determined under a luciferase multiple reader.

### Animal experiments

The SARS-CoV-2 infection of K18-hACE2 mice was conducted in compliance with the National Institutes of Health guidelines for the care and use of laboratory animals. Experiments were carried out in ABSL3 laboratories in accordance with the IACUC protocols of Texas Biomedical Research Institute (1718MU) and Cornell University (2017–0108).

### Comparison of the pathogenicity of BA.5, rBA.5, and rBA.5 NSP1ΔKSF *in vivo*

Five-month-old heterozygous male and female K18-hACE2 mice (strain: 2B6.Cg-Tg(K18- hACE2)2Prlmn/J) from The Jackson Laboratory were intranasally inoculated with the indicated inoculum of BA.5, rBA.5, or rBA.5 NSP1ΔKSF. Changes in body weight and survival were monitored daily. Mice reaching 25% body weight loss were humanely euthanized. Tissues from infected mice were collected for downstream analysis as indicated. In another experiment, mice were euthanized at 5 days post-infection (dpi), and brain, lung, and BALF samples were harvested to compare the virus load between different groups.

### Pathogenicity of rWA1 WT and rWA1 NSP1ΔKSF *in vivo*

Two-month-old female K18 hACE2 transgenic mice were intranasally infected (5×10^5^ PFU) with either rWA1 or rWA1 NSP1ΔKSF. After viral infection, three mice were randomly selected and humanely euthanized at 2 and 4 dpi, respectively, to collect nasal turbinate and lungs. The viral loads in nasal turbinate and lungs were determined as previously described (58). The rest of the mice (n=5) were used to assess changes in body weight and survival upon virus infection.

### Multiplex cytokine assay

After mice euthanasia, BALFs were obtained using 1 mL of PBS. Subsequently, centrifugation was utilized to remove cells for flow cytometry, and supernatants were processed on the ProcartaPlex™ Mouse High Sensitivity Panel, 5plex (ThermoFisher Scientific), following the manufacturer’s instructions. To neutralize SARS-CoV-2 present in BALFs, a 30-min inactivation step employing 10% formalin was used, followed by the addition of the Amplification Reagent and two consecutive washes. This was followed by a washing step before the final replacement with the Reading Buffer. Plates were analyzed using xPONENT software (Luminex) after reading on the Luminex MagPix instrument.

### Flow cytometry

Cells were collected from infected mice BALFs 5 dpi and stained and gated as previously described (17). Stained and fixed cells were run on the Attune NxT (Thermo Fisher Scientific) and analyzed using FlowJo Software 10 (BD Biosciences). Briefly, cells were gated on singlets and live cells before they were identified and graphed accordingly: alveolar macrophages (Ly6G-, CD11c+, SiglecF+), monocyte-derived macrophages (CD11b+F4/8 0+), NK cells (NK1.1+), neutrophils (Ly6G+, CD11b+), γδT cells (TCRγ+), NKT cells (TCRβ+, NK1.1+), CD4+ T cells (TCRβ+, CD4+), and CD8+ T cells (TCRβ+, CD8+).

### Histopathology and Immunolabeling

Tissues were collected at experimental endpoints and fixed in 10% formalin for at least 72 h before paraffin embedding. Tissue sections (4 μm) were stained by hematoxylin and eosin (HE) and blindly scored by an anatomic pathologist. The following criteria were used to assign scores based on the percentage of various tissue types (alveolus, vessels, etc.) affected: normal (0); less than 10% (1); between 10 to 25% (2); 26 to 50% (3); and higher than 50% (61). IHC sections were stained with an anti-SARS-CoV-2 N protein rabbit IgG monoclonal antibody (GeneTex; GTX635679) at a 1:5,000 dilution. Tissue sections were processed using a Leica Bond Max automated IHC stainer. Leica Bond Polymer Refined Detection (Leica; DS9800) with DAB as the chromogen. Images were acquired on a Roche Ventana DP200 slide scanner, and immunolabeling was blindly scored by an anatomic pathologist. The following criteria were used for scoring based on tissue affection: (0) none; (1) less than 10%; (2) 10-25%; (3) 26-50%; and (4) higher than 50%.

### Statistical Analysis

All statistical analyses were performed using GraphPad Prism v10.0. The primary methods included the Student’s t-test, Student’s t-test with Welch’s correction, Mann-Whitney test, and two-way ANOVA with appropriate multiple comparisons tests. For survival analysis, the log-rank (Mantel-Cox) test was applied. The specific statistical tests used for each experiment are indicated in the corresponding figure legends. Statistical significance is denoted as follows: *, p<0.05; **, p<0.01; ***, p<0.001; ****, p<0.0001; and non-significant differences are unmarked.

## ACKNOWLEDGMENTS

We thank the Cornell CARE staff for mouse colony maintenance and members of the Cornell BSL-3, ABSL-3, and Cornell Biosafety staff, including but not limited to P. Jenettee, N. Kushner, and J. Turse. We thank the Cornell Flow Cytometry Core, the Cornell Animal Health and Diagnostic Center Anatomic Pathology lab, and D. Gludish for comments and feedback. We also thank C. Florence at the NIH/NIAID and BEI Resources for sharing the SARS-CoV-2 Omicron BA.4 (NR- 56806) and BA.5 (NR-58620) strains. We also thank BEI Resources for providing Vero AT cells (NR- 54970).

Research in L.M-S.’s laboratory was partially funded, until March 24^th^, 2025, by the Antiviral Countermeasures Development Center, AC/DC (1U19AI171403-01); the Center for Antiviral Medicines & Pandemic Preparedness, CAMPP (1U19AI171443-01), and the QCRG Pandemic Response Program (1U19AI171110-01), three of the National Institutes of Health (NIH) funded Antiviral Drug Discovery Centers for Pathogens of Pandemic Concern; the Center for Research on Influenza Pathogenesis (CRIPT), the San Antonio Partnership for Precision Therapeutics, the San Antonio Medical Foundation, a Texas Biomedical Research Institute Forum Foundation to L.M-S., and a Texas Biomedical Research Institute Forum Foundation to C.Y.

Research in the Aguilar laboratory was partially funded by NIH grant R01AI109022 and Cornell University discretionary funds. This study was also supported by CAMPP, CRIPT, and NIAID grant R01AI184975 to A. G-S.

## AUTHORS CONTRIBUTIONS

C.Y., S.E., B.I., L.M-S., and H.C.A. conceived the study. C.Y., S.E., B.I., N.J., T.C., A. Cupic, L.M., A. Choi, D.W.B., J.C., J.S., X.A.O-C., M.C.J., and G.R.W. performed the experiments and analyzed the results. A.A., B.F., and A.G-S. provided critical reagents and methods. C.Y., S.E., and B.I. compiled the results. C.Y. and S.E. drafted the manuscript. L.M-S. and H.C.A. revised the manuscript. All authors reviewed and approved the final version.

---

**Supplemental Table 1.**
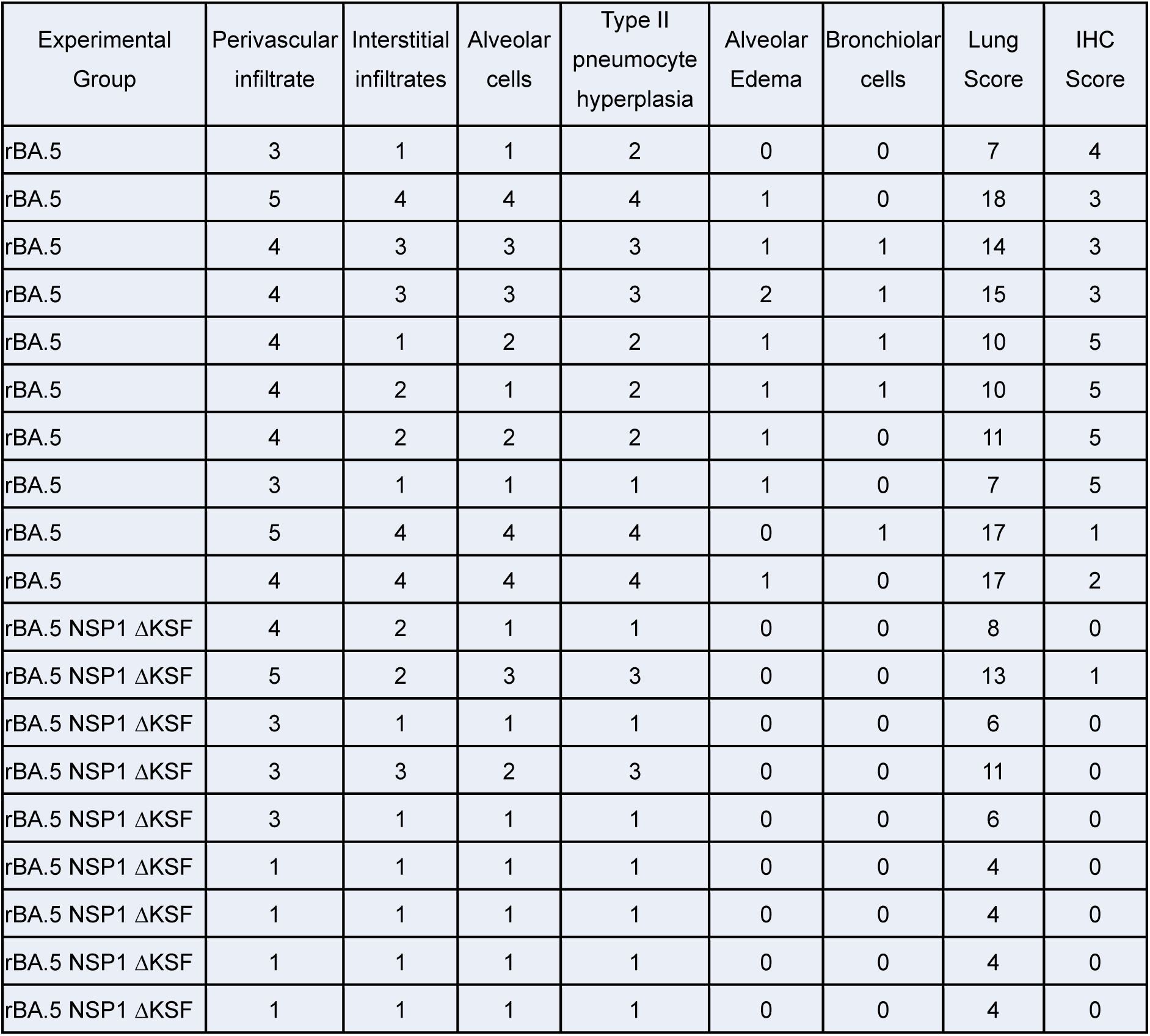
Histopathological and IHC Scoring of Lung at Humane or Study Endpoints: Lung tissue sections were evaluated for pathological changes by assigning scores based on the proportion of affected tissue, including alveoli, vasculature, and other relevant structures. The histological scoring criteria were as follows: 0 = no observable lesions (normal), 1 = <10% of the tissue affected, 2 = 10–25% affected, 3 = 26–50% affected, 4 = 51-75%, and 5 = >75% involvement. For IHC analysis, scoring was conducted on IHC-stained sections to assess antigen distribution and tissue involvement. The IHC scoring scale was defined as: 0 = no immunoreactivity (normal), 1 = <10% of the tissue showing positive staining, 2 = 10–25%, 3 = 26–50%, 4 = 51–75%, and 5 = >75% of the tissue affected. All evaluations were performed on formalin-fixed, paraffin- embedded sections, collected at designated time points post-infection.

| Experimental Group | Perivascular infiltrate | Interstitial infiltrates | Alveolar cells | Type II pneumocyte hyperplasia | Alveolar Edema | Bronchiolar cells | Lung Score | IHC Score |
| --- | --- | --- | --- | --- | --- | --- | --- | --- |
| rBA.5 | 3 | 1 | 1 | 2 | 0 | 0 | 7 | 4 |
| rBA.5 | 5 | 4 | 4 | 4 | 1 | 0 | 18 | 3 |
| rBA.5 | 4 | 3 | 3 | 3 | 1 | 1 | 14 | 3 |
| rBA.5 | 4 | 3 | 3 | 3 | 2 | 1 | 15 | 3 |
| rBA.5 | 4 | 1 | 2 | 2 | 1 | 1 | 10 | 5 |
| rBA.5 | 4 | 2 | 1 | 2 | 1 | 1 | 10 | 5 |
| rBA.5 | 4 | 2 | 2 | 2 | 1 | 0 | 11 | 5 |
| rBA.5 | 3 | 1 | 1 | 1 | 1 | 0 | 7 | 5 |
| rBA.5 | 5 | 4 | 4 | 4 | 0 | 1 | 17 | 1 |
| rBA.5 | 4 | 4 | 4 | 4 | 1 | 0 | 17 | 2 |
| rBA.5 NSP1 ΔKSF | 4 | 2 | 1 | 1 | 0 | 0 | 8 | 0 |
| rBA.5 NSP1 ΔKSF | 5 | 2 | 3 | 3 | 0 | 0 | 13 | 1 |
| rBA.5 NSP1 ΔKSF | 3 | 1 | 1 | 1 | 0 | 0 | 6 | 0 |
| rBA.5 NSP1 ΔKSF | 3 | 3 | 2 | 3 | 0 | 0 | 11 | 0 |
| rBA.5 NSP1 ΔKSF | 3 | 1 | 1 | 1 | 0 | 0 | 6 | 0 |
| rBA.5 NSP1 ΔKSF | 1 | 1 | 1 | 1 | 0 | 0 | 4 | 0 |
| rBA.5 NSP1 ΔKSF | 1 | 1 | 1 | 1 | 0 | 0 | 4 | 0 |
| rBA.5 NSP1 ΔKSF | 1 | 1 | 1 | 1 | 0 | 0 | 4 | 0 |
| rBA.5 NSP1 ΔKSF | 1 | 1 | 1 | 1 | 0 | 0 | 4 | 0 |

**Supplemental Table 2.**
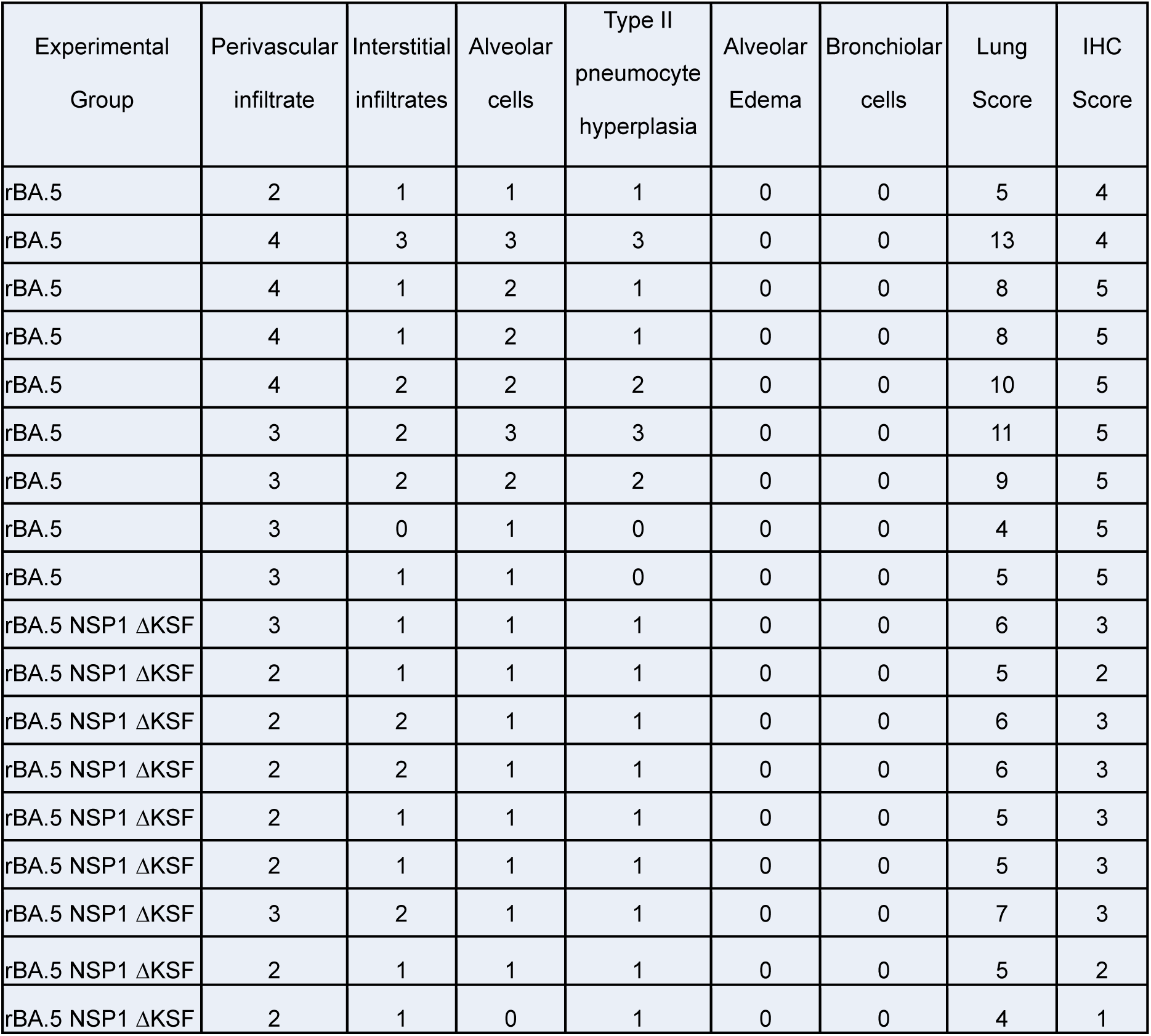
Histopathological and IHC Scoring of Lung at 5 dpi: Lung sections collected at 5 dpi were evaluated for pathology and antigen presence using previously described scoring criteria. Both histopathology and IHC scores ranged from 0 (normal) to 5 (>75% affected), based on the percentage of tissue involvement across key structures.

| Experimental Group | Perivascular infiltrate | Interstitial infiltrates | Alveolar cells | Type II pneumocyte hyperplasia | Alveolar Edema | Bronchiolar cells | Lung Score | IHC Score |
| --- | --- | --- | --- | --- | --- | --- | --- | --- |
| rBA.5 | 2 | 1 | 1 | 1 | 0 | 0 | 5 | 4 |
| rBA.5 | 4 | 3 | 3 | 3 | 0 | 0 | 13 | 4 |
| rBA.5 | 4 | 1 | 2 | 1 | 0 | 0 | 8 | 5 |
| rBA.5 | 4 | 1 | 2 | 1 | 0 | 0 | 8 | 5 |
| rBA.5 | 4 | 2 | 2 | 2 | 0 | 0 | 10 | 5 |
| rBA.5 | 3 | 2 | 3 | 3 | 0 | 0 | 11 | 5 |
| rBA.5 | 3 | 2 | 2 | 2 | 0 | 0 | 9 | 5 |
| rBA.5 | 3 | 0 | 1 | 0 | 0 | 0 | 4 | 5 |
| rBA.5 | 3 | 1 | 1 | 0 | 0 | 0 | 5 | 5 |
| rBA.5 NSP1 ΔKSF | 3 | 1 | 1 | 1 | 0 | 0 | 6 | 3 |
| rBA.5 NSP1 ΔKSF | 2 | 1 | 1 | 1 | 0 | 0 | 5 | 2 |
| rBA.5 NSP1 ΔKSF | 2 | 2 | 1 | 1 | 0 | 0 | 6 | 3 |
| rBA.5 NSP1 ΔKSF | 2 | 2 | 1 | 1 | 0 | 0 | 6 | 3 |
| rBA.5 NSP1 ΔKSF | 2 | 1 | 1 | 1 | 0 | 0 | 5 | 3 |
| rBA.5 NSP1 ΔKSF | 2 | 1 | 1 | 1 | 0 | 0 | 5 | 3 |
| rBA.5 NSP1 ΔKSF | 3 | 2 | 1 | 1 | 0 | 0 | 7 | 3 |
| rBA.5 NSP1 ΔKSF | 2 | 1 | 1 | 1 | 0 | 0 | 5 | 2 |
| rBA.5 NSP1 ΔKSF | 2 | 1 | 0 | 1 | 0 | 0 | 4 | 1 |

**Supplemental Figure 1.**
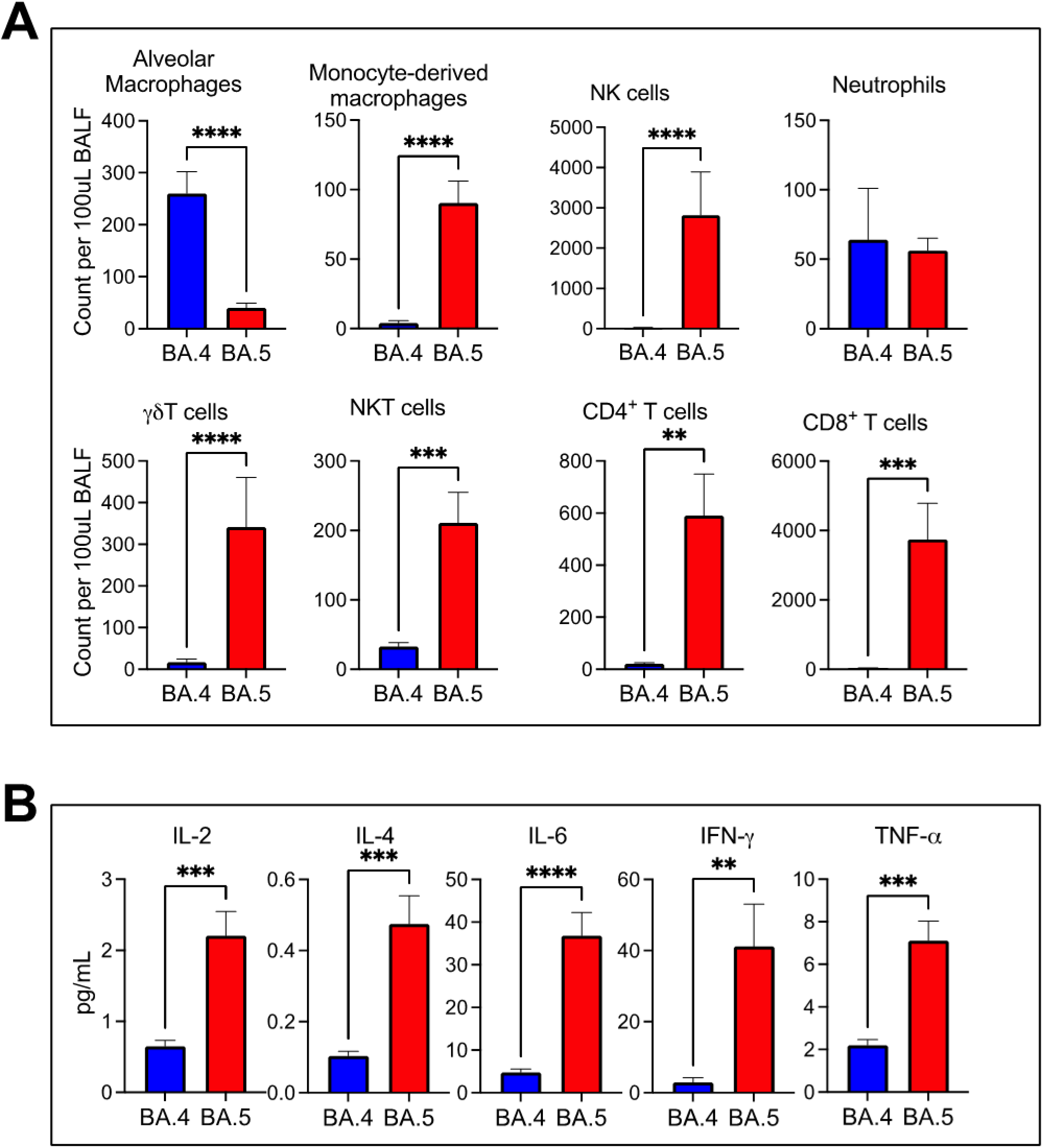
Immune cells and cytokines in BALF from K18-hACE2 mice infected with BA.5 and BA.4: BALF were collected from K18-hACE2 mice (5-8 months old, n=10: 5 male and 5 female) with the infection (3.25×10^4^ PFU/mouse) of BA.5 and BA.4 at 5 dpi and analyzed by flow cytometry (**A**). The clarified supernatant of BALF was used to measure the levels of the indicated cytokines using multiplex (**B**). Data are presented as Mean ± SD. Statistical significance was determined using unpaired Student’s *t*-tests with Welch’s correction. **, *p*<0.01; ***, *p*<0.001; ****, *p*<0.0001.

**Supplemental Figure 2.**
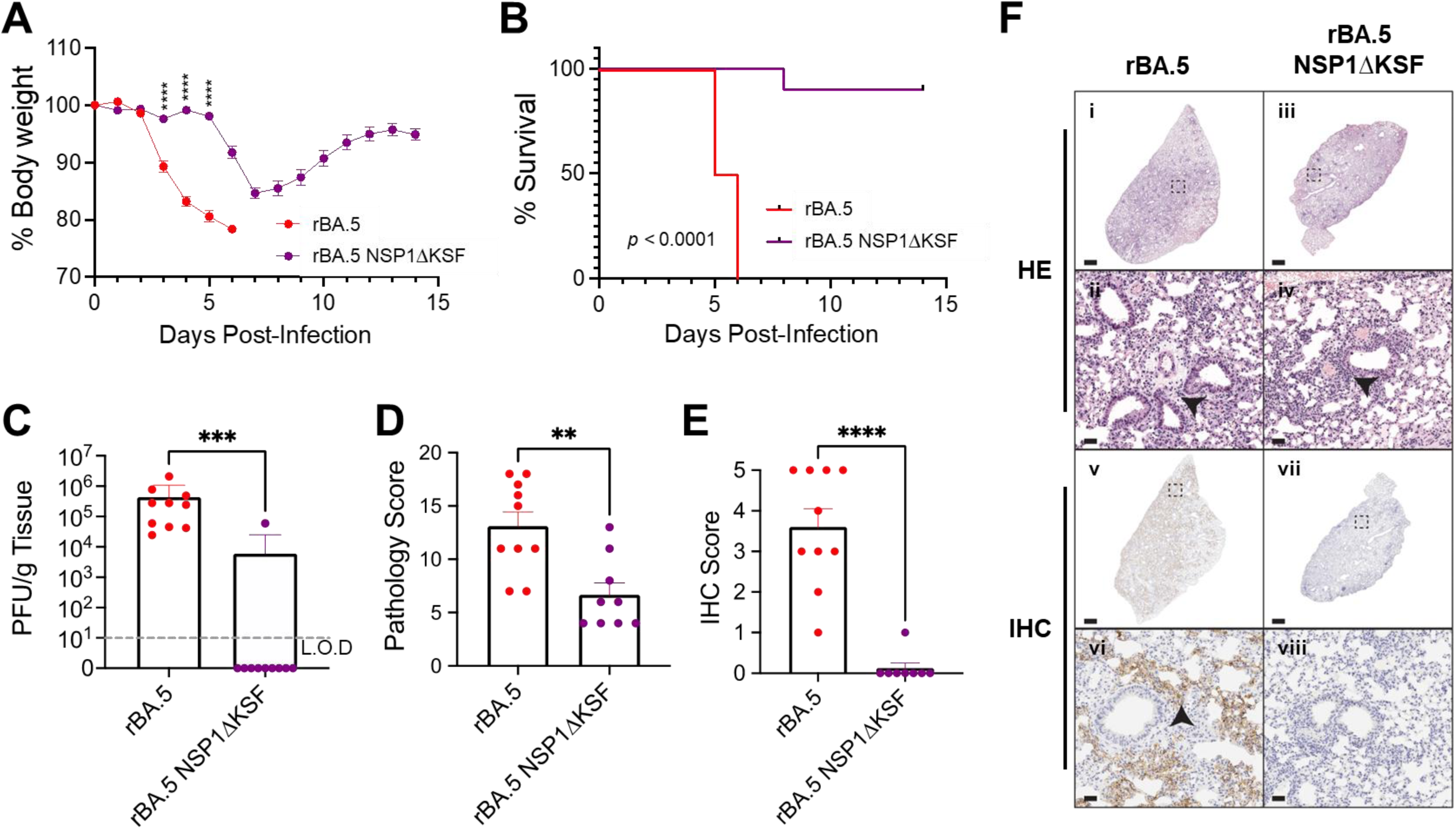
The triple KSF deletion in NSP1 reduced BA. 5-induced disease severity *in vivo*: K18--hACE2 (5-8 months old) mice were infected intranasally with 1×10^5^ PFU/mouse of rBA.5 or rBA.5 NSP1ΔKSF (n=10: 5 male and 5 female per group). Change in body weight was monitored daily for 14 days (**A**), and survival was plotted accordingly (**B**). The lungs were collected at humane or experimental endpoints for viral titers (**C**), pathology analysis (**D**), and IHC staining (**E**). (**F**) Lung pathology (i–iv) and antigen distribution (v–viii) in rBA.5- and rBA.5 NSP1ΔKSF-infected mice. Perivascular (arrowheads in ii and iv) inflammatory infiltrates, along with viral antigen presence (arrowhead in vi), were more prominent in lungs from rBA.5-infected mice compared to the NSP1ΔKSF group. Scale bars: 1 mm (i, iii, v, vii); 100 μm (ii, iv, vi, viii). Images are representative of group averages based on pathology and IHC scoring at humane or experimental endpoints. Data are presented as Mean ± SEM. Survival differences between the groups of rBA.5 and rBA.5 NSP1ΔKSF were analyzed using the log-rank (Mantel-Cox) test. Unpaired Student’s *t*- tests with Welch’s correction were used to compare the means of the titers. Mann-Whitney tests were used to compare the differences between groups for the scores of pathology and IHC. **, *p*<0.01; ***, *p*<0.00; ****, *p*<0.0001.

